# Influence of neural network bursts on functional development

**DOI:** 10.1101/2025.01.21.633559

**Authors:** Ola Huse Ramstad, Axel Sandvig, Stefano Nichele, Ioanna Sandvig

## Abstract

1

Network bursts or synchronized burst events are a typical activity seen in most in vitro neural networks. Network bursts arise early in development and as networks mature, activity becomes dominated by bursts propagating across the entirety of the network. The reason for this developmental plateau in vitro is unknown, but to bypass it would confer a significant advantage in the use of in vitro networks for computation. As most neurons in a network participate in network bursts, burst onset supersedes any ongoing activity thereby placing a limit on short term computations equal to the recovery period between bursts. By assessing 521 multielectrode array recordings from day 4-39 in vitro we find that network bursts influence the connectivity, but this change is only weakly associated with the origin of the network bursts. The impacts of bursts on functional integration and segregation in neural networks are discussed along with approaches to mitigate the development and propagation of network bursts in vitro. Additionally, we hypothesize that burst initiation zones or pacemakers are viable targets for stimulation for computation in the context of control and reservoir.

**2#Author summary:** When neurons from the cortex are grown in culture outside the brain, they begin to recapitulate some of the initial activity seen during development. The activity is initially sporadic but over time it becomes more driven by bursts of activity which often stem from only a few areas or neurons in the culture. These bursts of activity often come to make up most of the activity in the culture, making it highly synchronized but also reducing the variety of activity. In this article, we examine whether the areas from which the bursts originate drive the formation of connections between neurons as the culture develops, therefore making signals propagate more effectively to the rest of the network. To that end we analysed the recorded activity from 30 cultures as they developed over 39 days and found that while these areas do influence the development of connections between the neurons, they do not propagate their signals more effectively over time.

## 3 Introduction

### 3.1 Background

Neural networks in vitro almost invariably trend towards the propagation of network bursts (1–7). These network bursts, also termed population bursts (8) or synchronized burst events, are a common feature of maturation in dissociated in vitro networks across multiple cell types and experimental conditions (3,6,9–12). Multiple hypotheses persist as to the origin of these network bursts including coupling of intrinsic oscillations (6), calcium transients (1,13), hypometabolic responses (14), topological features, (15,16) and pacemaker neurons (2,10,13,16–19). As dissociated neurons develop in vitro, mechanisms of self-organization and Hebbian plasticity increase both synchrony and the propagation of network bursts to the point where mature networks frequently contain most spiking activity within these population events (13,20,21). This prevalence of network bursting can thus dominate the activity of the in vitro network, pushing out more varied and complex activity (22–24), and in turn reduce its information capacity in terms of short-term computations (25–27).

A better understanding of network bursts and, thereby, our ability to control them can significantly improve the validity of in vitro neural networks as computational substrates (23). As others have posited (3,20,23,28), this form of continuous bursting activity represents a key step in the normal development of cortical neurons in vivo by creating an increased synchrony between the neurons in the network. However, the lack of thalamic input, or indeed any external input to drive the in vitro networks beyond this activity is a challenge. Arguably the only input provided to developing in vitro neural networks is mechanical perturbation during media change and recording. Thus, the network burst persistence likely represents a form of arrested development, though the cause of this developmental plateau is still unknown.

### 3.2 Problem

We posit that the network bursts, originating from pacemakers or burst initiation zones (2), act as the main drivers of the network’s functional development. Ultimately, this will be represented in the connectivity as an increasing efficiency towards propagation of signals from these pacemakers, decreasing the pathlength between this and all other nodes in the network. To show this we have utilized all dense network recordings provided by the Wagenaar, Pine & Potter dataset (3) to track both the initiation of network bursts, their stability and how the connectivity or propagation pattern of each network burst stabilizes to ensure this synchrony and ease of propagation. We have detected each network burst in the recordings, its origin point, spatiotemporal propagation pattern, the within and between recording stability of these patterns, and compared these to the developing functional connectivity and network metrics. We discuss identifying and selectively targeting such pacemakers during network development to enable more complex behaviour that better reflects the functional dynamics of in vivo networks.

## 4 Results

### 4.1 Burst development and stability

As described, in vitro networks trend towards increased network bursting as they mature. By assessing the burstiness index (figure 1b) across DIV a clear trend (r = 0.62, p < 0.0001) towards increased bursting is evident across almost all batches. Yet as noted by (3) there is a more consistent trend within batches than between, with some batches (1,5,8) showing mostly burst confined spikes and others (3) only moderate increases. In addition to the increase in network burst prevalence the duration between spikes within bursts increases across development. By plotting the ISI between every spike t and t+10 we can see the shift in both burst contained and tonic firing across development (figure 1a). As the network matures the ISI within bursts decrease, as do ISI non-burst spikes.

**Figure 1:**
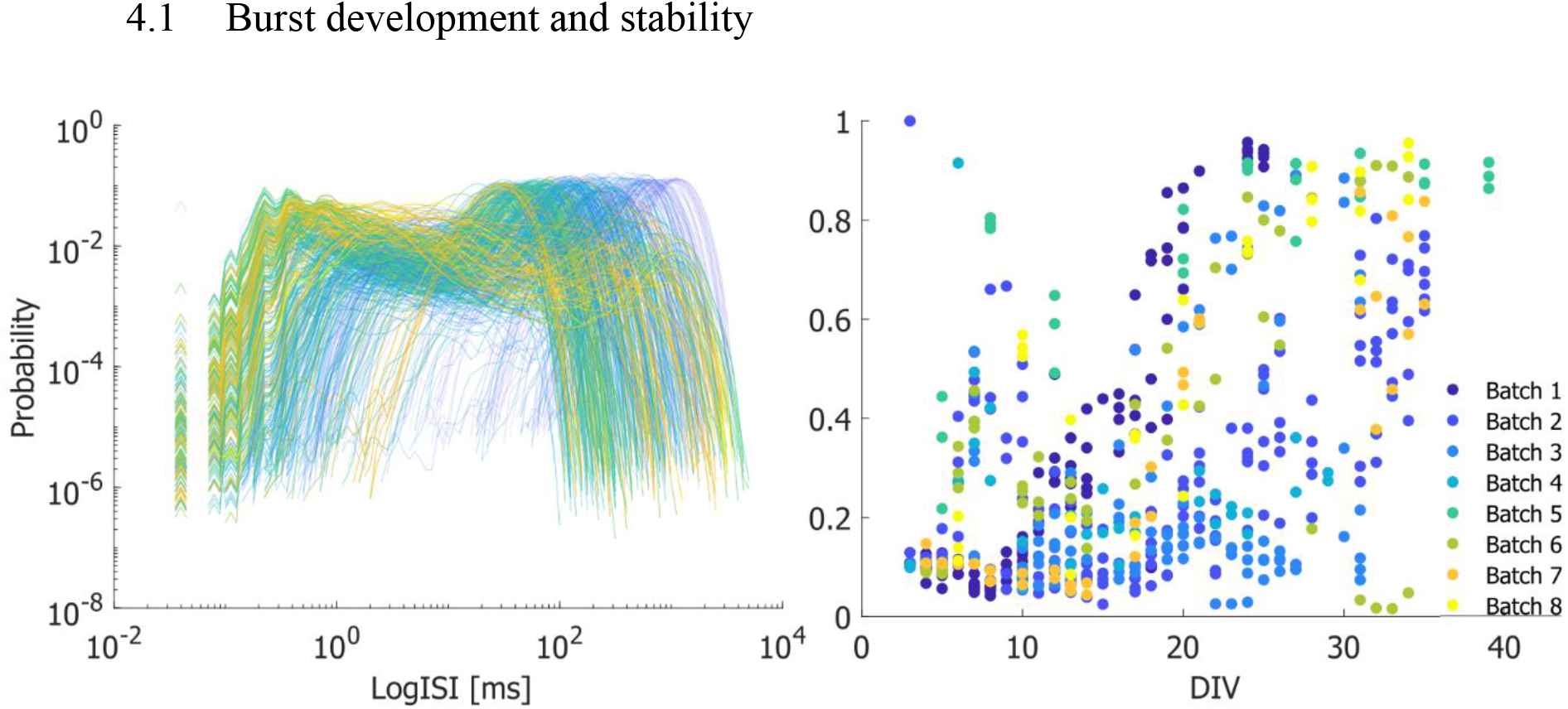
LogISI plot and burstiness index across network development. (a) Early burst ISI (blue) show a peak (right) at longer intervals than late bursts (yellow) indicating that burst spiking becomes more rapid as the network matures. (b) The burstiness of the networks also increase with DIV (r = 0.62, p < 0.0001) across all batches and MEAs.

This reduction in burst ISI does not necessarily mean that the durations of the network bursts themselves decrease at the same rate (figure 2c). The duration of network bursts is highly variable in the initial onset period between DIV 5-14 before they stabilize on brief durations between DIV 15-20 then gradually increase in length from DIV 20 onwards. These trends are also highly batch dependent with some MEAs showing short and stable network bursts from their onset.

**Figure 2:**
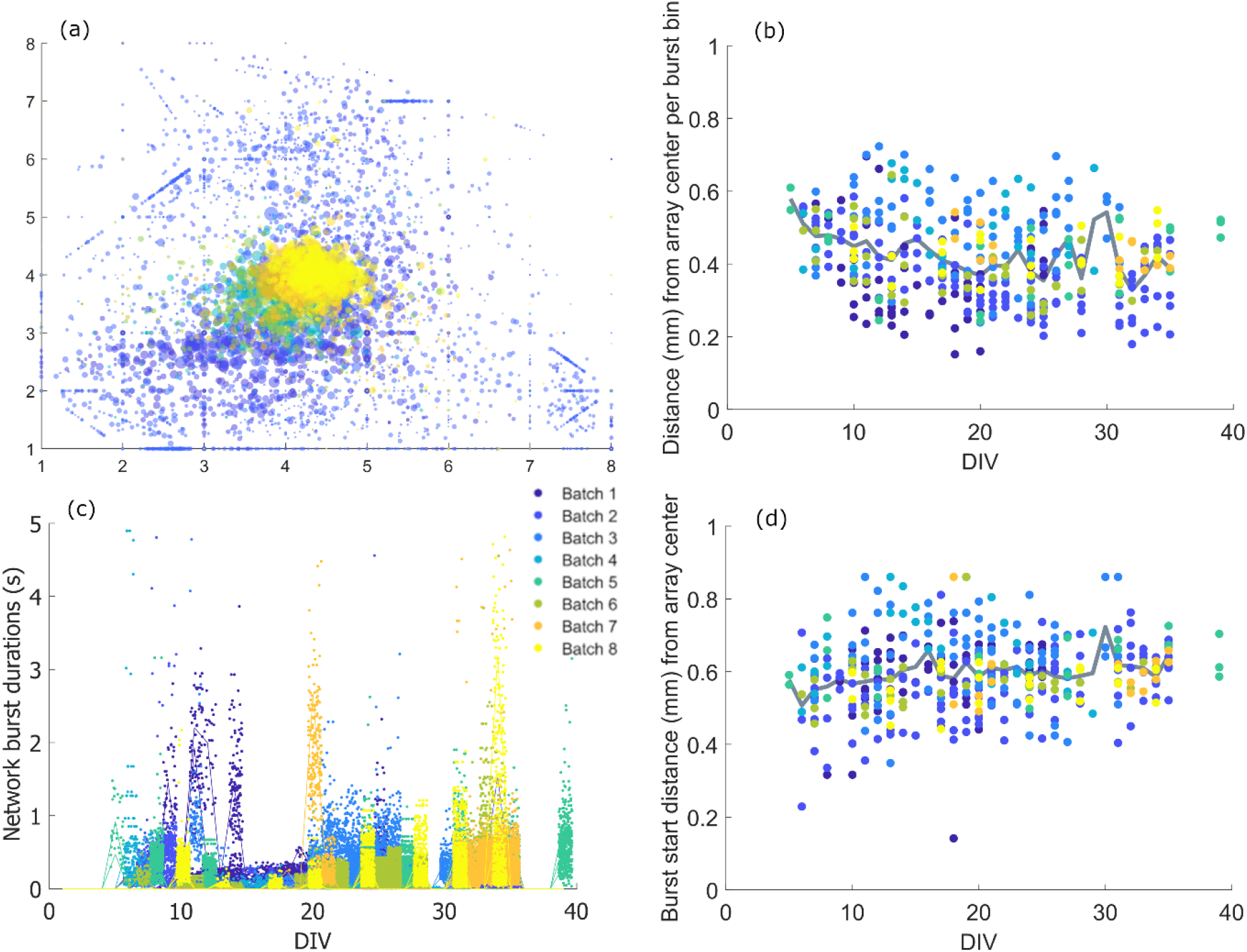
Center of burst activity and stability over time. (a) Burst centers ordered from early (blue) to late (yellow) DIV. Marker size given by total spike count per 10ms burst frame. (b) As network bursts increase in spike count and electrode recruitment the burst centers weakly shift from peripheral to central locations on the array as seen by the decreasing mean distance from center (r = -0.25, p < 0.0001, mean shown in grey). (c) Network burst duration varies across MEA batches and time. Early (5–13) DIV network bursts show varied durations and shapes compared to later DIV though this differs between batches. (d) The starting electrodes of the network bursts are located on the outlying electrodes with a weak shift away from the central electrodes as the networks mature (r = 0.26, p < 0.0001, mean shown in grey).

While spiking activity in general is evenly distributed across the electrode array, the bursting activity and especially network bursting is dependent on the network age (figure 2a). Initially the center of activity during bursting is confined to the edge of the array, near pacemakers or the initial electrode activated during a detected network burst. These centers of activity gradually shift (r = -0.25, p < 0.0001) towards the middle of the array as more electrodes are recruited during each network burst and the average spiking is more evenly distributed across the array (figure 2b). The initial electrode activated during network bursts stays consistent at the edge of the array (figure 2d) with a small change away from the center across the maturation of the network (r = 0.26, p < 0.0001). The reason for this is likely twofold, as network bursts originate outside the array they will first be detected at the edge, secondly, persistent paths develop along already active electrodes. The difference in pacemaker and non-pacemaker-initiated network bursts were not significantly different (figure 3a) and burst alignment did not change across DIV (figure 3b, r = -0.04, p = 0.44).

**Figure 3:**
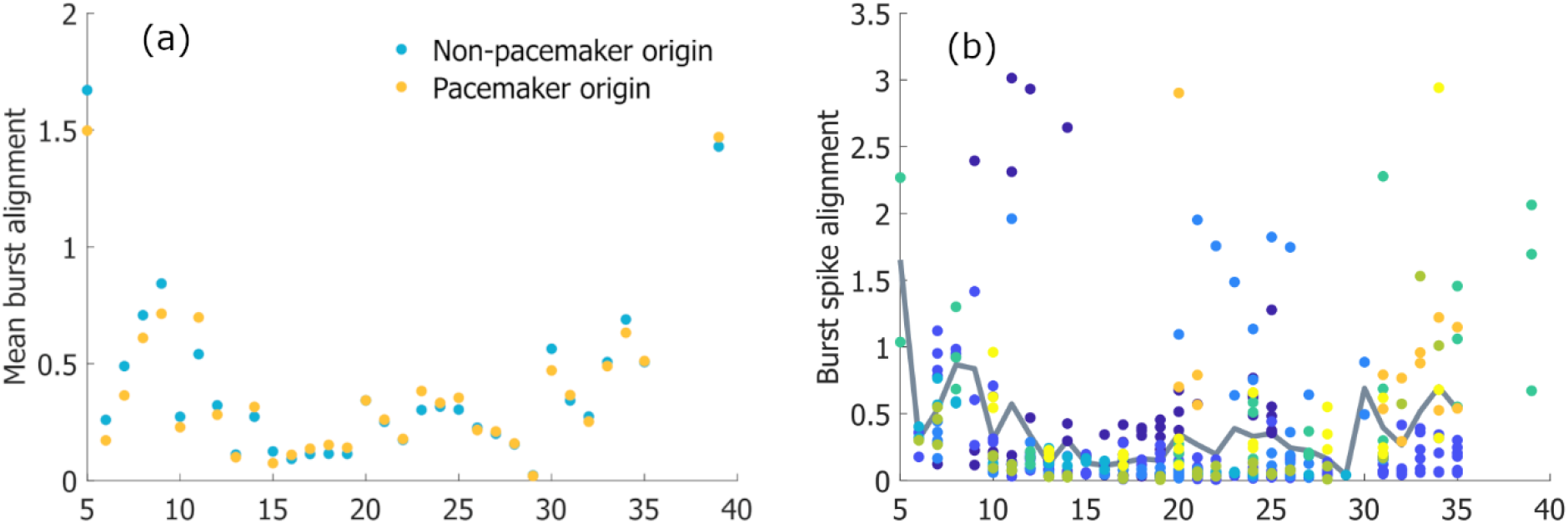
Network burst alignment across DIV. (a) The alignment of time to first spike network burst patterns per day are not significantly different in bursts originating from pacemakers. (b) The network burst alignment is dependent on batch, with some batches trending towards increasing alignment across DIV (lower distance per day) while some remain stable. Overall burst alignment did not change across DIV (r = -0.04, p = 0.44). Mean alignment across all batches shown in grey.

### 4.2 Impact of pacemakers on network development

Lag time pathlengths from pacemakers are marginally higher across development, only approaching the network median after DIV 30 (figure 4a). The betweenness centrality of the pacemakers does not deviate from the network median (figure 4b) however, the degree centrality (figure 4c) is lower than that of the giant component from 8 DIV to 21 DIV. The mutual information and clustering coefficient increase from 8 DIV (figure 4d-e) for the pacemakers and remain consistently higher than the giant component median. The modularity (figure 4f) started high early in development before decreasing between 5-10 DIV and slowly decreasing from 10 DIV onwards. Overall, the lag time metrics for the pacemakers did not deviate significantly from the giant component, while mutual information metrics for the pacemakers showed some deviation from 16 DIV (Supplementary figure 1). There was a significant increase in mutual information (r = 0.52, p < 0.0001), clustering (r = 0.52, p < 0.0001) and BI (r = 0.62, p < 0.0001) over DIV, however there is only a weak correlation between mutual information and BI (r = 0.18, p < 0.0001). Conversely there is a decrease in lag path length (r = -0.18, p < 0.0001), betweenness (r = -0.27, p < 0.0001) and degree centrality (r = -0.48, p < 0.0001) over DIV. The decrease in modularity and increase in BI show a strong negative correlation (r = -0.63, p < 0.0001), contrasting the relationship with mutual information and BI.

**Figure 4:**
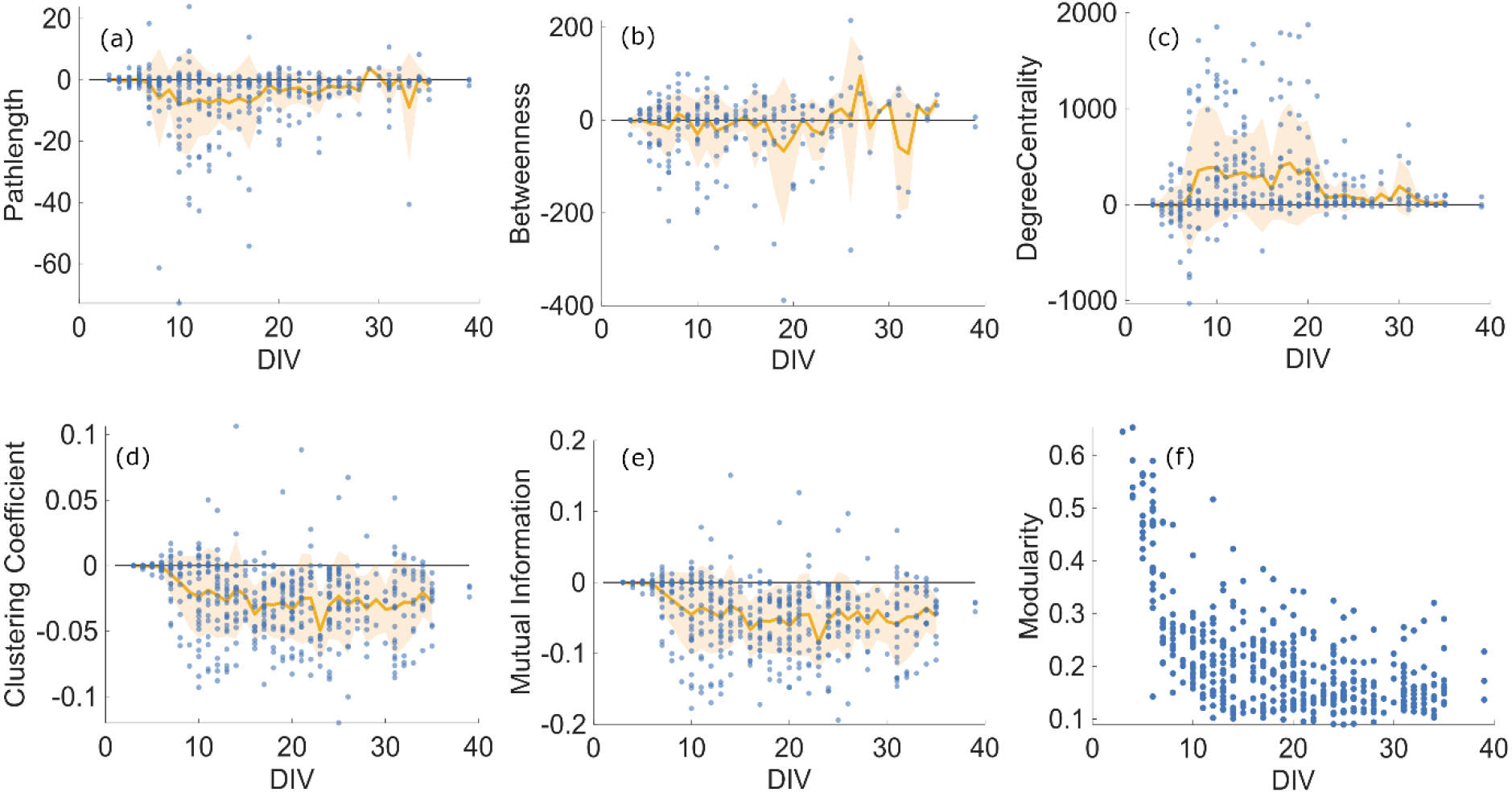
Comparison of network characteristics between pacemakers and giant component. a-c calculated using lag times, d-f calculated using mutual information. Plots a-e show difference in median between pacemaker and giant component per MEA per day in blue. Orange line denotes mean difference with shaded standard deviation. Pathlengths (a) between pacemakers and other network nodes are marginally longer on average than random nodes 8-30 DIV. Betweenness (b) on average shows no difference except for some variability from 26-35 DIV. Degree (c) centrality is lower than the network mean consistently from 8-20 DIV but approaches the network mean from 20 DIV onwards. Clustering (d) and mutual information (e) is consistently higher for the pacemakers from 8 DIV onwards. Modularity (f) for the giant component begins high but rapidly declines till 10 DIV and slightly decreases from 10 DIV onward.

## 5 Discussion

We hypothesized that network bursts, originating from areas of the network called pacemakers or burst initiation zones, act as a main driver of the in vitro network’s functional development. To investigate this hypothesis, we utilized dense network recordings provided by the Wagenaar dataset to track the initiation, stability, and connectivity of network bursts. As the network develops, the efficiency of signal propagation increases, resulting in increasing mutual information yet this only weakly corresponded with the BI. The pathlengths of pacemakers were consistently higher than the network mean, however pacemakers were not central in the network as per betweenness and degree centrality. Pacemakers followed the trend of the giant component in reducing path lengths and increasing mutual information of DIV while retaining lower degree centrality from 8-20 DIV.

Considering that bursting networks show increasing levels of integration, network bursting may be enabling information to propagate more effectively across the network. However, without sufficient segregation, typically in the form of hierarchical modularity, this high integration does not lend itself to an increase in input separability. In this study we found a rapid initial decrease from high to low modularity before the q stabilized. Given the modularity maximization of the Louvain method, small size of the network, and homogeneity of the culture vessel it is possible this high initial modularity stemmed from spurious modules of a few interconnected electrodes which shifted to more stable modules which encompass the majority of the array. On larger MEAs and CMOS arrays, Okujeni and Egert (2,13,20) demonstrated well the increased segregation which arises from physical neural aggregation, with subsequent increases in variability of activity and reduced recruitment from network bursts. The issue with continuous network bursts is twofold in terms of computation (34), firstly, the increase in synchrony reduces the network segregation and eventual diversity of activity. Secondly, the onset of a network burst typically means ongoing spiking activity is superseded by the network burst, thus putting a limit on short term computations, including fading memory, equivalent to the length of interburst intervals. Note that the term computation is often non-specific and context dependent, we use the term here specifically to denote the use of *in vitro* networks for applied computational tasks, such as performance in robot/animat control (35–38), or in terms of physical reservoir computing (39–42).

To increase the computational viability of in vitro culture it is necessary to find approaches that reduce the long term impact of network bursts (23). Though certain neurons in a network are likely more intrinsically active (10), it is probable that the network burst initiation seen in vitro arises from self-organization as detailed by Penn et al. (43) and Lonardoni et al. (44). Disrupting the development of network bursts therefore would mean disrupting the self-organization itself, either through perturbing neurite growth (20), establishing compartments (45,46), complex cell composition (47,48) or altering the inhibitory-excitatory balance (49). In our previous work we have impeded the propagation of network bursts by the addition of global GABAergic inhibition to the cultures (50), though this method is transient due to the rapid uptake and recycling of free GABA. A more long-term solution based on the excitation-inhibition ratio will entail altering the prevalence of interneurons or their connectivity in the networks (48). Future studies may benefit from further exploring the role of the excitation-inhibition ratio in this context (51).

Most studies on MEAs using dissociated neurons perform recordings in the period 14 to 40 DIV, regardless of species of origin or cell type. Synchronizing population bursting also occurs in vivo in rodents during late embryonic and early postnatal cortical development before typically declining as non-local input increases (52,53). The timescale for this initiation and decline of population bursting therefore occurs on shorter timescales than the typical recording period of MEA studies. Yet as reductionist models, dissociated cultures are lacking key features of neural development including glial cell and extracellular matrix (ECM) composition, three-dimensional structure, and gradient compartmentalization. Induced pluripotent stem cells (IPSCs) do recapture some of these factors in terms of their cell composition and matrix complexity, but more importantly studies on IPSCs frequently extend across months and up to a year in length (47,54,55). While IPSCs do show a similar trend in network bursting early in development, some IPSC networks eventually stabilize and decline in burstiness depending on the differentiation protocol in contrast to rat cortical neurons which can sustain bursting up to two years (3,56). Extending recording periods beyond the standard 30-40 DIV may therefore capture more variable activity depending on the cell type.

In studies utilizing in vitro networks for computations, stimulations or inputs to the networks are usually applied to random electrodes to reduce bias, or central electrodes to capture as much of the resulting response as possible(36,38,56,57). Some studies improve upon this by selecting more active electrodes, yet the network structure itself is rarely considered in terms of input and output selection (36,58). Given that neural networks in vitro self-organize to propagate network bursts, either due to developmental set points or as a function of culture conditions, more care should be taken to both optimize integration and segregation of inputs by considering the network structure. Utilizing pacemakers as input points while concurrently increasing the heterogeneity and complexity of in vitro networks should enable better computational capacity than is currently achievable with these networks.

## 6 Materials and methods

### 6.1 Dataset

All data was obtained from Wagenaar et al., (2006), please see original article for details on experimental setup and spike detection. For this analysis, spike times from all *dense* (seeding density of 2.5 ± 1.5*10^3^ cells per mm^2^) recordings of the dataset (n=521) were selected. All analysis was performed using MATLAB (R2025b, Mathworks). As detailed in the original article, recordings were affected by artefacts due to movement from the incubator to the recording stage. To remove the artefacts and reduce noise in the burst detection, the initial 200 seconds of each recording were removed prior to analysis. Further, brief recordings (<5 minutes following artefact removal) were excluded to keep recordings of comparable length. In total, recordings from 30 microelectrode arrays (MEAs) across 8 batches were included, spanning DIV 3-39.

### 6.2 Burst detection

Bursts were detected on a per electrode basis using the log interspike interval (logISI) method detailed by (29). In brief, using a base 10 logISI histograms, bursts can be detected using adaptive interspike interval (ISI) thresholding as spikes in bursts typically display shorter ISI from those not contained in bursts. This divergence can be used to find a recording specific maximum ISI for burst detection as the last local minima (or 100ms if exceeded) on a logISI histograms and a minimum of 4 spikes within each interval. For burst centers, given a 100ms bin, the center of mass/activity is the weighted sum of firing rates for each neuron in the array. The distance between burst centers and the center of the array is given by the Euclidean distance of the cartesian coordinates of the center of mass and the coordinates [4.5, 4.5] in the 8x8 array.

### 6.3 Network burst detection

For network burst detection an adaptation was made to this method. Based on the bursts detected, a vector of burst times was generated for each recording. As each recording displayed a different burst length and profile which consistently changed across DIV, an adaptive binning for the network burst detection was used. Bursts were binned using the *mode* of the burst durations creating a bursts per bin vector. Taking the derivative of the bursts per bin allows the detection of network bursts as the positive peaks above the median burst rate or 3, whichever is higher. This number was selected to correspond to a minimum number of electrodes necessary for a network burst as well as differentiate those recordings with a high non-network burst rate. To find the start and end time of each network burst, the first bin before and after the peak with a derivative burst rate below the median was selected. As network bursts at early (10–14) DIV and late DIV (25+) display greatly differing burst durations and shapes, this adaptive approach was necessary to detect and compare multiple types.

### 6.4 Burstiness index and pacemaker detection

Burstiness index (BI) was calculated with a population (all electrodes) spike binning of bin size 1s sorted in order of highest to lowest spike count and calculating fraction of spikes contained in the top 15% (f_15_) of the bins. This is then normalized as (f_15_-0.15)/0.85 (23). Finally, pacemakers or burst initiation zones were selected as follows: the first electrode to burst following the start time of a detected network burst as noted, disregarding spurious electrodes with less than 2 activations per recording. Then we summed the number of times each electrode initiated a network burst across all recordings per MEA and defined pacemakers as those electrodes initiating at least 4% of network bursts (18,30).

### 6.5 Similarity of network bursts

For network burst alignment we created a vector of the time delay between the start of a network burst and the first spike which occurs at each electrode. That is, given a network burst lasting from time t(s) to t(e) what is the time delay to the first spike in this duration for each electrode. Time delays exceeding 1 second were discarded to better ensure spikes were related to the onset of the network burst. For each network burst a 1x59 vector of time delays was used to assess similarity. Once vectors of time delays are given for each network burst the similarity can be estimated giving a similarity matrix across all bursts. For all network bursts recorded per MEA the Euclidian distance between network bursts was estimated with missing values accounted for by using the root of squared distances scaled by the ratio of all electrodes to valid pairs.

### 6.1 Functional connectivity

Functional connectivity was estimated using cross-correlation and mutual information. First, spikes were binned in 100ms bins, and an initial correlation was estimated using Spearman’s rank correlation then filtered to discard correlations with p > 0.01 and r < 0.1. Remaining correlations were assessed using normalized cross-correlation with a bin size of 1ms and a maximum lag of ±100ms. Optimal lag times were selected based on the maximum correlation within the tested delays. Mutual information was calculated using 100ms bins, and each potential connection was tested for significance by comparing the initial mutual information to the mutual information of a randomly shuffled binning for the same connection. This shuffling was performed 100 times with connections deemed significant if the initial mutual information exceeded 95% of the shuffled cases. In comparing the effect of network burst propagation on network formation, the pacemakers were first detected as described in section 3.4. Using the detected pacemakers, the contribution of these nodes to the network development could be described.

### 6.2 Network metrics

The characteristic path length is defined as the mean of the shortest path length between all pairwise nodes, where the distance between nodes i and j is defined as 1/w_ij_ using the optimal lag time. Betweenness and degree centrality were calculated using the path lengths as described. Clustering and modularity were estimated using mutual information. The clustering coefficient of a node was defined as the ratio of the number of connections between the node’s neighbors to the maximum number of possible connections between them. The local coefficient of each node is calculated using the method by Onnela et al. (31) for weighted networks (Brain Connectivity toolbox, BCT). Modularity q was estimated using the Louvain method (32) as implemented in the BCT. For each network the modularity was calculated 100 times to account for variability in the community detection, with the q being given as the median of the 20 estimates. Network metrics are shown as the difference of the median of the giant component minus the median of the pacemakers. Correlations between development (DIV), BI, and network metrics were performed with repeated measures correlation (33) to account for within-MEA associations across time. Correlation p-values below 0.0001 are reported as 0.0001.

## Code availability

All code is available on GitHub under MIT License following appropriate citation: https://github.com/Olahr-brain/Network-burst-connectivity

## Data availability

All original data is available on https://neurodatasharing.bme.gatech.edu/development-data/html/index.html

## Acknowledgements

Funding was provided by the DeepCA project (Research Council of Norway, Young Research Talent grant agreement 286558).

## Author contributions

OHR: conceptualization, methodology, data and statistical analysis, writing. SN, AS, IS: conceptualization, writing, review, and editing. All authors contributed to the article and approved the submitted version.

## 8 Supplementary

### 8.1 Bootstrap distributions of network metrics

Violin plots were generated using bootstrapping of the difference scores between pacemakers and giant component network metrics. For each DIV with at least n=8 scores for that network metric, a bootstrapping was performed by taking n samples with replacement 1000 times. The means of each bootstrap sampling were then used to generate the distributions and 95% confidence intervals for each violin plot. As the underlying metric of the bootstrap is a measure of difference, the deviation from 0 is interpreted as significant, specifically where the 95% confidence intervals do not include 0.

**Supplementary figure 1:**
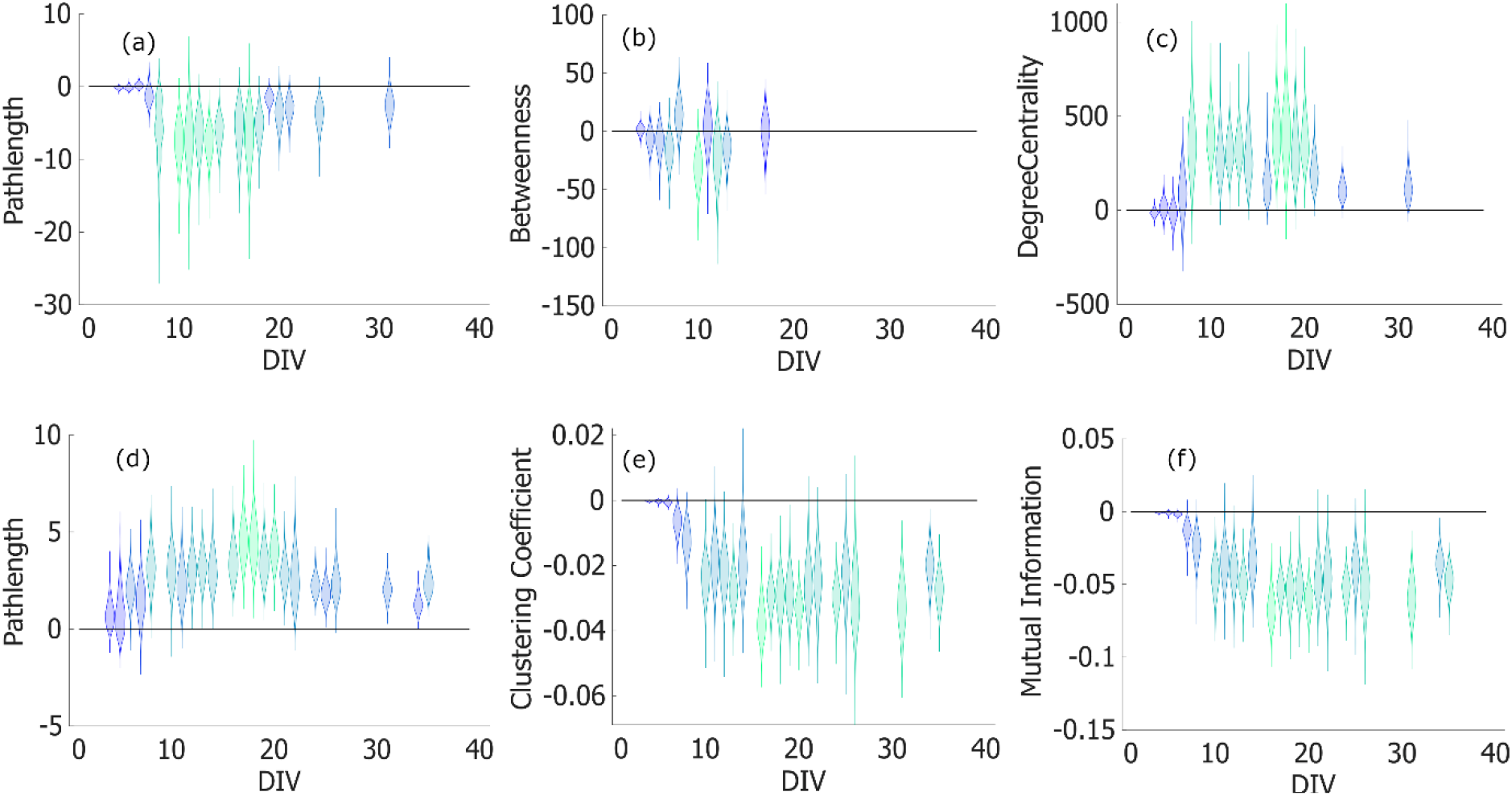
Violin plots showing bootstrap distributions of network characteristics from difference between pacemakers and giant component. a-c calculated using lag times, d-f calculated using mutual information. Colour denotes distance of the distribution mean from 0, with lighter colours showing greater deviation. Lag time metrics for the pacemakers showed no, or little significant deviation from the giant component. Mutual information metrics show some significant deviation between 16 DIV and 21 DIV and from 30 DIV onwards.

